# *Bdnf* promoter IV-expressing cells in the hippocampus modulate fear expression and hippocampal-prefrontal synchrony in mice

**DOI:** 10.1101/552646

**Authors:** Henry L. Hallock, Henry M. Quillian, Yishan Mai, Kristen R. Maynard, Julia L. Hill, Keri Martinowich

## Abstract

Brain-derived neurotrophic factor (BDNF) signaling regulates synaptic plasticity in the hippocampus (HC) and prefrontal cortex (PFC), and has been extensively linked with fear memory expression in rodents. Notably, disrupting BDNF production from promoter IV-derived transcripts enhances fear expression in mice, and decreases fear-associated HC-PFC synchrony, suggesting that *Bdnf* transcription from promoter IV plays a key role in HC-PFC function during fear memory retrieval. To understand how promoter IV-derived BDNF affects fear expression and HC-PFC connectivity, we generated a viral construct that selectively targets cells expressing promoter IV-derived *Bdnf* transcripts (“p4-cells”) for tamoxifen-inducible Cre-mediated recombination (AAV8-p4Bdnf-ER^T2^CreER^T2^-PEST). Using this construct, we found that ventral (vHC) p4-cells are recruited during fear expression, and that activation of these cells causes exaggerated fear expression that co-occurs with disrupted vHC-PFC synchrony in mice. Our data highlight how this novel construct can be used to interrogate genetically-defined cell types that selectively contribute to BDNF-dependent behaviors.

## Introduction

Fear dysregulation is a core symptom of several neuropsychiatric disorders (Orr & Roth, 2000; Lommen et al., 2013). In rodents, freezing (cessation of movement) to a fear-conditioned cue or context (e.g., a tone or environment that has been paired with a foot-shock) is routinely used as a proxy for fear memory. Fear expression in rodents depends on several brain areas, including the HC (Kjelstrup et al., 2002; Trivedi and Coover, 2004), mPFC (Morgan and LeDoux, 1995), and amygdala (Anglada-Figueroa and Quirk, 2005; Ciocchi et al., 2010). The rodent HC routes information to both the prelimbic (PrL) and infralimbic (IL) subregions of the mPFC (Swanson, 1981; Jay and Witter, 1991), which contribute to fear expression and extinction, respectively (Sierra-Mercado et al., 2011). The HC and mPFC interact during contextually-mediated fear expression and extinction (Lesting et al., 2011; Hill et al., 2016), suggesting that vHC-mPFC communication is critical for fear memory.

vHC-mPFC synapses are plastic (Jay et al., 1996), suggesting that structural changes in this circuit underlie its ability to support fear memory. Brain-derived neurotrophic factor (BDNF) is a signaling molecule that regulates synaptic plasticity during development and in the adult brain (Akaneya et al., 1997; Lu, 2003), and many studies demonstrate a relationship between *BDNF* gene expression and fear memory in rodents (Bredy et al., 2007; Lubin et al., 2008; Soliman et al., 2010). Infusion of BDNF protein in the HC-IL circuit facilitates fear extinction (Peters et al., 2010), suggesting that BDNF regulates fear behavior by controlling plasticity in the hippocampal-prefrontal network (Hill & Martinowich, 2016).

*BDNF* is transcribed from several unique promoters, which are activated by different signaling cascades to initiate promoter-specific transcription (Timmusk et al., 1993, West et al., 2001; Aid et al., 2007; West et al., 2014). Transcription from *Bdnf* promoter IV (p4) is activity-dependent: exon IV-containing *Bdnf* transcripts are upregulated following membrane depolarization (Shieh et al., 1998) and kainic acid-induced seizures (Metsis et al., 1993). Recent evidence suggests that BDNF production from individual promoters differentially impacts fear memory; specifically, BDNF disruption from promoter IV (p4)-derived transcripts impairs fear expression in mice (Sakata et al., 2013), and decreases fear-related hippocampal-prefrontal oscillatory synchrony (Hill et al., 2016). These results suggest that cells expressing BDNF from promoter IV (“p4-cells”) critically regulate hippocampal-prefrontal plasticity during fear memory.

To selectively tag “p4-cells” for labeling and manipulation, we developed an adeno-associated adenovirus (AAV) expressing an ER^T2^CreER^T2^ fusion protein under control of *Bdnf* promoter IV. Utilization of this construct with 4-hydroxy-tamoxifen (4OHT) causes indelible expression of Cre recombinase only in p4-cells. We leveraged this construct to selectively label and manipulate p4-cells in the vHC of behaving mice during recall of both a fear-associated context and a fear-associated tone/context combination, allowing us to causally probe relationships between activity of these cells, fear-related behavior, and fear-related vHC-mPFC circuit dynamics. We found that vHC p4-cells are recruited during learned fear expression, and that synthetic activation of these cells increases fear expression while attenuating fear-related oscillatory synchrony both within the vHC and between the vHC and mPFC.

## Results and Discussion

We first cloned a tamoxifen-inducible Cre-recombinase expression cassette (ER^T2^CreER^T2^-PEST) downstream of the proximal mouse *Bdnf* promoter IV sequence (initial 600 bases upstream of the exon IV transcription start site), which was then inserted into a pAAV1 backbone and packaged as high titer adeno-associated virus (AAV8-p4BDNF-ER^T2^CreER^T2^-PEST; Cyagen Biosciences). To validate that the p4BDNF-ER^T2^CreER^T2^ construct undergoes recombination selectively in the presence of 4OHT, we co-injected AAV8-p4BDNF-ER^T2^CreER^T2^-PEST with Cre-dependent AAV8-CAG-FLEX-tdTomato into the vHC of adult mice (Fig. 1a). Two weeks later, we administered 4-hydroxy-tamoxifen (4OHT) for three consecutive days (20 mg/kg, i.p.) for Cre/lox-mediated expression of tdTomato protein. Two weeks following 4OHT injection, animals were killed, and tdTomato expression was compared between vehicle (saline) and 4OHT-injected mice. Robust tdTomato expression was observed in the ventral dentate gyrus of 4OHT-injected (Cre+/4OHT+) mice, while virtually no tdTomato expression was seen in saline-injected (Cre+/4OHT-) mice (Fig. 1c). We found virtually no tdTomato expression in mice injected with AAV8-FLEX-tdTomato and 4OHT in the absence of p4BDNF-ER^T2^CreER^T2^ (Cre-/4OHT+; Fig. 1c), demonstrating that Cre-mediated recombination requires the presence of 4OHT (Indra et al., 1999). In Cre+/4OHT+ mice, anti-BDNF immunohistochemistry revealed that a majority (∼95%) of tdTomato-expressing cells co-expressed BDNF protein (Supp. Fig. 1a). To investigate whether *Cre* and *tdTomato* transcripts were co-expressed with p4-derived *Bdnf* transcripts, we used single-molecule fluorescence *in situ* hybridization (RNAscope) to visualize virally-induced *Cre, tdTomato,* and exon IV-containing *Bdnf* transcripts in individual vHC neurons (Colliva et al., 2018; Fig. 1d). *Cre* transcripts were expressed only in Cre+/4OHT+ and Cre+/4OHT-animals (Fig. 1e), while *tdTomato* mRNA expression was limited to Cre+/4OHT+ mice (Fig. 1f), further demonstrating that 4OHT allows Cre to flip and excise the FLEX cassette for *tdTomato* transcription and translation. In Cre+/4OHT+ mice, the number of exon IV-containing *Bdnf* transcripts was significantly higher in *Cre*+ as compared to *Cre*-cells, and *tdTomato*+ compared to *tdTomato*-cells (Fig. 1g). We trained a binary classifier using logistic regression to determine whether the number of exon IV-containing *Bdnf* transcripts could predict *tdTomato* co-expression, and found that the classifier performed with high accuracy compared with shuffled data (Fig. 1h-i), demonstrating that the p4BDNF-ER^T2^CreER^T2^ construct targets p4-cells with high precision.

**Figure 1.**
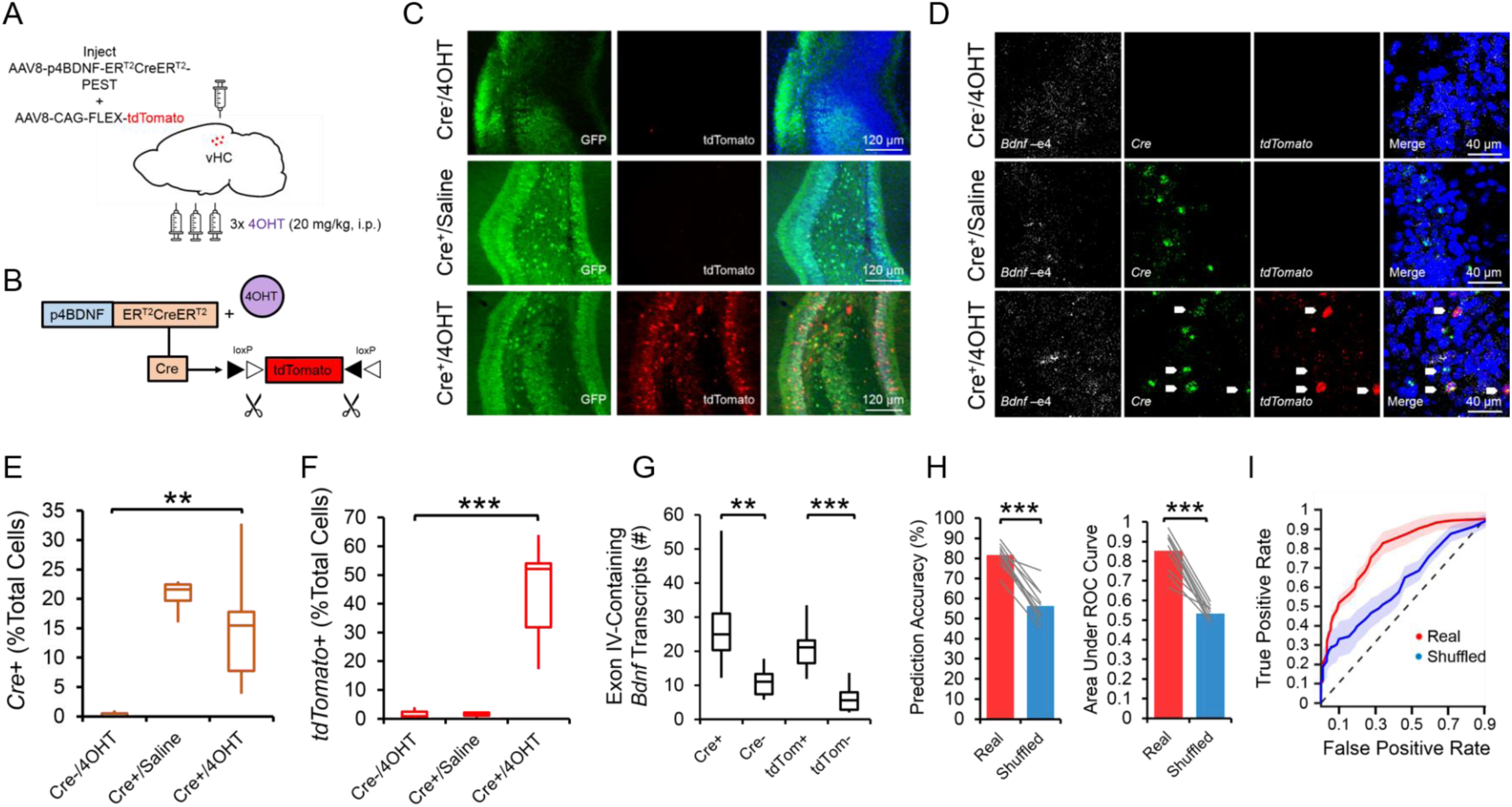
The p4BDNF-ER^T2^CreER^T2^ construct selectively targets cells expressing exon IV-containing *Bdnf* transcripts in the vHC (p4-cells). *(A)* Schematic of injection strategy for labeling of p4-cells in the vHC. *(B)* 4OHT binds to the ER^T2^CreER^T2^ fusion protein to allow Cre/lox-mediated expression of tdTomato protein. *(C)* tdTomato protein expression is observed only in animals that received injections of p4BDNF-ER^T2^CreER^T2^ and 4OHT (GFP labeling from co-injections of AAV1-hSyn-GFP were used to verify injection accuracy). *(D)* Single-molecule *in situ* hybridization reveals expression of *Cre* mRNA in animals that received injections of p4BDNF-ER^T2^CreER^T2^, and *tdTomato* expression is restricted to animals that received injections of both p4BDNF-ER^T2^CreER^T2^ and 4OHT. *(E)* A significantly higher percentage of cells co-express *Cre* in animals that received injections of p4BDNF-ER^T2^CreER^T2^, in comparison to animals that received injections of 4OHT without p4BDNF-ER^T2^CreER^T2^ *(F*(2,15) = 7.72, *p* = 0.005, one-way ANOVA). *(F)* A significantly higher percentage of cells co-express *tdTomato* mRNA in animals that received both p4BDNF-ER^T2^CreER^T2^ and 4OHT, as compared to both control groups *(F*(2,15) = 25.35, *p* < 0.0001, one-way ANOVA)*. (G)* Cells that express either *Cre* or *tdTomato* co-express a higher number of exon IV-containing *Bdnf* transcripts, as compared to cells that do not express *Cre (t*(9) = 4.44, *p* = 0.0016, paired t-test) or *tdTomato (t*(9) = 6.66, *p* < 0.0001, paired t-test)*. (H)* A logistic regression model trained to predict whether a cell expresses *tdTomato* based on number of exon IV-containing *Bdnf* transcripts performs with high accuracy for observed data, but not data in which number of *Bdnf* transcripts is randomly shuffled relative to *tdTomato* expression probability (*t*(9) = 9.82, *p* < 0.0001, paired t-test; left panel). Receiver operating characteristic (ROC) curves on classifier performance confirm model accuracy in real and shuffled data (*t*(9) = 9.67, *p* < 0.0001, paired t-test; right panel). *(I)* Regression classifier ROC curves for real and shuffled data (dashed line = chance).

To investigate whether p4-cells are recruited during fear expression, we again labeled ventral dentate gyrus p4-cells with tdTomato, and compared levels of the immediate early gene product c-Fos in tdTomato-labeled p4-cells between mice that received a tone-shock combination during conditioning, and mice that received only the tone (no shock; Fig. 2a). Shocked mice froze at significantly higher levels than no shock mice during conditioning, context recall, and tone/context recall (Fig. 2b). Mice were killed 2 h after tone/context recall to measure c-Fos levels during fear expression. A higher percentage of p4-cells co-expressed c-Fos in the shock group compared to the no shock group (Fig. 2d), while total number of c-Fos+ cells did not significantly differ between conditions (Supp. Fig. 2a). We next asked whether synthetic excitation of p4-cells is sufficient to potentiate fear expression during recall. To answer this question, we expressed the excitatory DREADD receptor hM3Dq in p4-cells by co-infusing AAV8-p4BDNF-ER^T2^CreER^T2^-PEST with AAV8-hSyn-DIO-hM3Dq-mCherry in the vHC (“hM3Dq” group). 45 minutes before context recall, we administered clozapine-N-oxide (CNO, 5 mg/kg, i.p.) to engage the hM3Dq receptor in p4-cells (Fig. 2e). Activation of vHC p4-cells caused increased freezing during both context and tone/context recall, as compared to CNO-injected animals that expressed a control reporter (AAV8-hSyn-DIO-mCherry, “mCherry” group; Fig. 2f). Effects of vHC p4-cell activation on fear expression were not due to general deficits in movement or anxiety, as neither locomotor (Supp. Fig. 2b-c) nor anxiety-like behavior (Supp. Fig. 2d-f) differed between groups post-CNO injection. Ensuring that CNO activated vHC p4-cells, the percentage of mCherry cells co-expressing c-Fos was significantly higher in hM3Dq mice following CNO injections (Fig. 2h). These results demonstrate that vHC p4-cells represent a sub-population capable of enhancing contextually-mediated fear memory recall.

**Figure 2.**
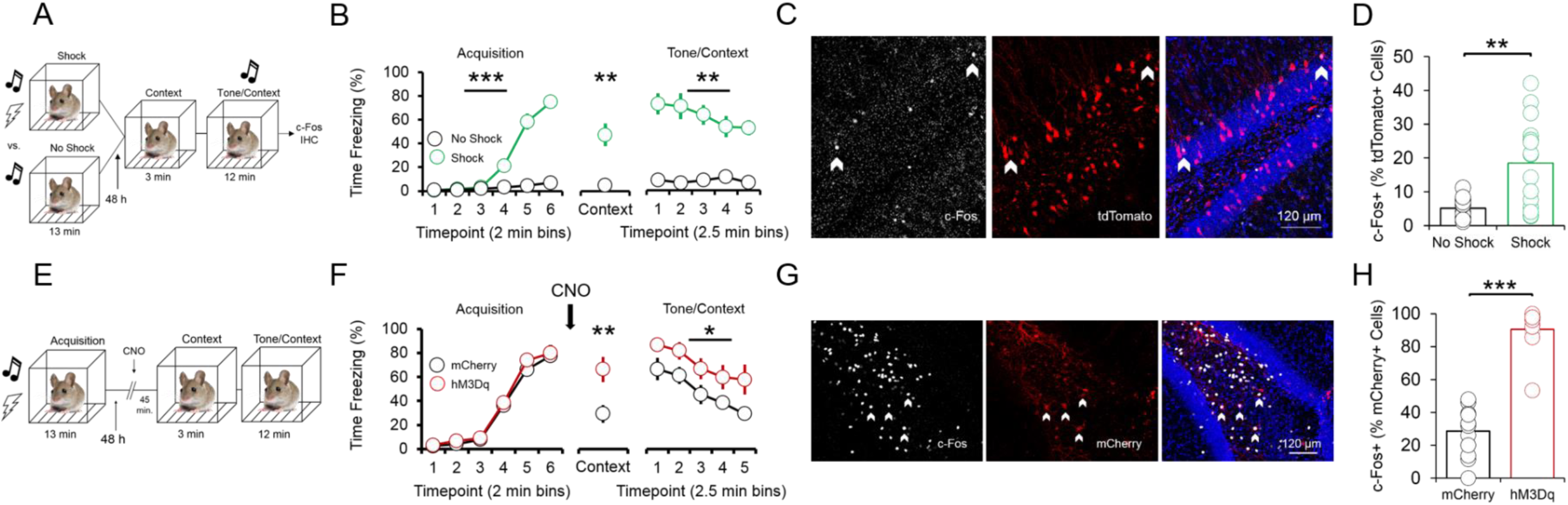
p4-cells in the vHC enhance fear expression during context and tone/context recall. *(A)* Schematic of experimental design for assessing recruitment of vHC p4-cells during fear expression. All animals received injections of p4BDNF-ER^T2^CreER^T2^, FLEX-tdTomato, and 4OHT prior to fear conditioning. Half of the animals received a tone-shock combination during conditioning, and the other half received only the tone. *(B)* Animals in the shock group froze significantly more frequently than animals in the no shock group during conditioning (*F*(1,5) = 73.58, *p* < 0.0001, group x time interaction), context recall (*t*(10) = 4.37, *p* = 0.0014, unpaired t-test), and tone/context recall (*F*(1,4) = 5.4, *p* = 0.0014, group x time interaction). *(C)* c-Fos immunohistochemistry was used to identify p4-cells in the vHC that were recruited during recall. *(D)* A higher proportion of p4-cells (tdTomato+) in the vHC co-express fear-induced c-Fos in mice that received a shock during conditioning (*t*(14) = 4.1, *p* = 0.0003, unpaired t-test). *(E)* Schematic of experimental design for assessing whether vHC p4-cells enhance fear expression during recall. All animals received injections of p4BDNF-ER^T2^CreER^T2^ and 4OHT, and half of the animals additionally received injections of an AAV encoding for a Cre-dependent excitatory DREADD (hM3Dq) fused to a fluorescent reporter (mCherry), while the other half received injections of an AAV coding for the Cre-dependent expression of only mCherry. *(F)* Synthetic activation of vHC p4-cells prior to context recall significantly increases freezing during both context recall (*t*(18) = 2.96, *p* = 0.01, unpaired t-test) and tone/context recall (*F*(1,18) = 6.08, *p* = 0.02, main effect of condition). *(G)* c-Fos immunohistochemistry was used to verify that CNO injections activate hM3Dq-expressing vHC cells. *(H)* A significantly higher proportion of mCherry+ cells in the vHC co-express c-Fos following CNO injections in mice that express hM3Dq-mCherry, as compared to mice that express only mCherry (*t*(24) = 11.75, *p* < 0.0001, unpaired t-test).

Systemic reductions in p4-derived BDNF cause fear expression deficits that co-occur with decreased hippocampal-prefrontal synchrony (Hill et al., 2016), suggesting that p4-cells modulate vHC-mPFC circuit function to impact fear behavior. To test this hypothesis, we recorded simultaneous local field potentials (LFPs) from the vHC and PrL, a sub-region of the PFC that is highly implicated in fear expression, during context and tone/context recall in hM3Dq and mCherry mice after CNO injections. Activation of vHC p4-cells did not significantly affect broadband power in the vHC (Fig. 3b-d); we further found that slow gamma oscillations (30-50 Hz) preferentially coupled to theta oscillations (4-12 Hz) within the vHC (Fig. 3e) at higher-than-chance levels (Supp. Fig. 3a) during tone/context recall (Fig. 3f-g), and that theta-gamma coupling in the vHC was significantly lower in hM3Dq mice following CNO injections (Fig. 3h). Decreased theta-gamma coupling in the vHC was concomitant with decreased theta phase coherence between the vHC and PrL (Fig. 3i-j), suggesting that activation of vHC p4-cells disrupts vHC-PrL synchrony to promote fear expression. Disruptions in vHC-PrL theta synchrony were directionally-specific, as the vHC strongly led the PrL in all frequency bands measured in mCherry mice; this directionality was selectively disrupted in the theta frequency band in hM3Dq mice (Fig. 3k) during tone/context recall (Fig. 3l). CNO injections did not significantly affect theta-gamma coupling (Supp. Fig. 3c), or theta phase coherence (Supp. Fig. 3f) in the home cage.

**Figure 3.**
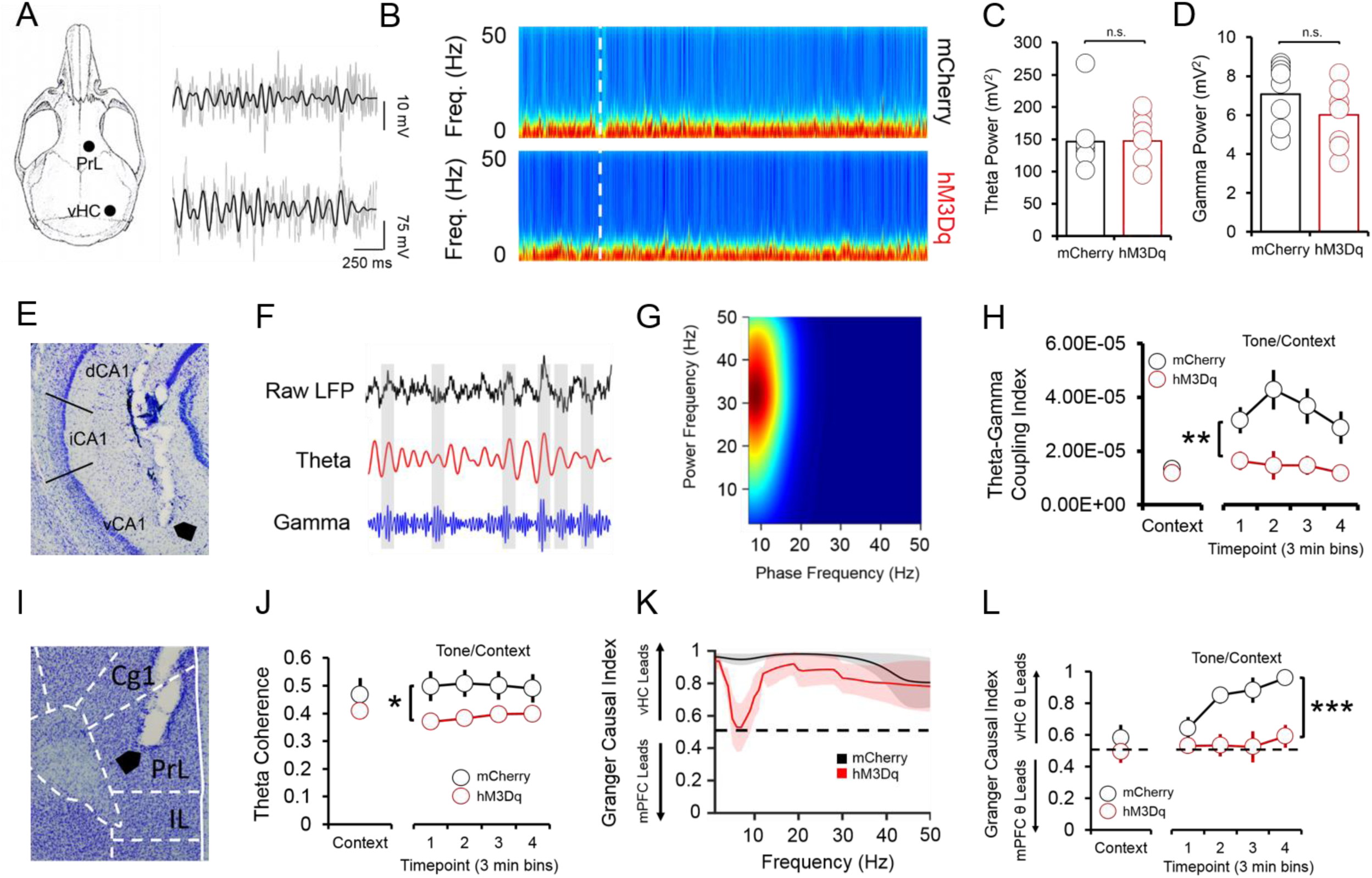
Fear-related vHC-PrL synchrony is disrupted following activation of p4-cells in the vHC. *(A)* Schematic of electrode placements and sample LFP recordings from the PrL and vHC (grey), with filtered theta superimposed on top (black). *(B)* Spectrograms from the vHC in hM3Dq (hM3Dq-mCherry in p4-cells) and mCherry (mCherry only in p4-cells) mice during fear recall. White line = start of tone/context recall. *(C)* Neither theta (4-12 Hz) power, nor *(D)* slow gamma (30-50 Hz) power significantly differ between hM3Dq and mCherry mice during context and tone/context recall (*p* > 0.05 for individual samples t-tests). *(E)* Histology showing placement of an electrode tip in the vHC. *(F)* Within the vHC, bouts of high-amplitude slow gamma oscillations tended to occur around the peaks of the theta oscillation during fear recall. *(G)* This phase-amplitude coupling occurred most strongly between hippocampal theta (phase on the x-axis) and slow gamma (amplitude on the y-axis) oscillations (heat = coupling index). *(H)* Theta-gamma coupling was significantly lower during tone/context recall in mCherry mice (*F*(1,17) = 10.71, *p* = < 0.01, main effect of condition). *(I)* Histology showing an electrode tip in the PrL. *(J)* Phase coherence between theta oscillations in the vHC and PrL was significantly lower in mCherry mice during tone/context recall (*F*(1,17) = 8.34, *p* = 0.01, main effect of condition). *(K)* The vHC LFP led the PrL LFP in all frequency bands during tone/context recall in mCherry mice, while CNO injections severely disrupted this effect selectively in the theta frequency band in hM3Dq mice (black dashed line = chance). *(L)* vHC theta strongly led PrL theta during tone/context recall in mCherry mice, and this effect was abolished in hM3Dq mice (*F*(1,17) = 28.57, *p* < 0.0001, main effect of condition).

Our results describe a novel method for targeting genetically-distinct cell populations based on promoter-specific BDNF expression, expanding on previous research using the CreER^T2^ system to label and manipulate immediate early gene (IEG)-expressing neurons during behavior (Denny et al., 2014; Ye et al., 2016). We used this method to demonstrate that p4-cells cells in the ventral dentate gyrus form a sub-population that increases learned fear expression. Increased fear expression following activation of these cells co-occurred with decreased vHC-PrL synchrony, suggesting that these neurons regulate fear expression by dampening vHC-PrL communication. These results provide insights into how sub-populations of cells defined by isoform-specific expression of *Bdnf* can regulate complex behavior and circuit function, and contribute to a growing understanding of how BDNF signaling impacts hippocampal-prefrontal function in fear-related behavior.

## Materials and methods

### Animals

Male wild-type (w/t) C57Bl6/J mice were group-housed (3-5 animals per cage) and maintained on a 12 h light/dark cycle in a temperature and humidity-controlled colony room. All animals had *ad libitum* access to food and water. Animals were 10-16 weeks of age, and weighed 25-35 g at time of surgery. All procedures were in accordance with the SoBran Institutional Animal Care and Use Committee.

### Surgeries and 4OHT injections

Mice were deeply anesthetized with isoflurane (1-2.5% in oxygen), placed into a stereotaxic frame, and an incision was made along the midline of the scalp. The skull was leveled, bregma was identified, and small holes were drilled with a 0.9mm burr (Fine Science Tools) above the vHC (−3.2mm AP, ±3.1mm ML) for viral injections. For all injections, AAV8-p4BDNF-ER^T2^CreER^T2^ was first diluted in filtered 1x PBS at a ratio of 1:4 (for animals used for immunohistochemistry and *in situ* hybridization) or 1:6 (for animals used for behavior/electrophysiology). For immunohistochemistry and *in situ* hybridization experiments, 5 µl of pre-diluted AAV8-p4BDNF-ER^T2^CreER^T2^ was mixed with 5 µl of AAV8-CAG-FLEX-tdTomato (Addgene) and 5 µl of AAV1-hSyn-eGFP-WPRE-bGH (Addgene; also pre-diluted to a 1:4 ratio in filtered 1x PBS), for a total dilution ratio of 1:12 AAV8-p4BDNF-ER^T2^CreER^T2^, 1:3 FLEX-tdTomato, and 1:12 hSyn-eGFP. For behavior/electrophysiology experiments, 5 µl of pre-diluted AAV8-p4BDNF-ER^T2^CreER^T2^ was mixed with 5 µl of either AAV8-hSyn-DIO-mCherry (control group) or AAV8-hSyn-DIO-hM3Dq-mCherry (experimental group) for a total dilution ratio of 1:12 AAV8-p4BDNF-ER^T2^CreER^T2^ and 1:2 hM3Dq/mCherry. A total volume of 600 nl/hemisphere was injected into the vHC (−3.4mm DV) at a rate of 200 nl/min via a 10 µl syringe (Hamilton) controlled by an automated infusion pump (World Precision Instruments). For LFP recordings, two stereotrodes (35 µm stainless steel, California Fine Wire) were also implanted into the vHC and mPFC (+2.0 AP, ±0.3 ML, −1.8 DV). Stereotrodes were attached to a headmount, along with two wires soldered to two bone screws (Fine Science Tools). Ground screws were implanted above the frontal cortex on the hemisphere opposite the mPFC electrode, and directly above the lambda skull suture. Mice received a sub-cutaneous injection of a local anesthetic (Bupivicaine) along the incision site, as well as an i.p. injection of an analgesic (Meloxicam, 5 mg/kg) during surgery. Meloxicam injections were given for three days post-surgery for pain relief. Two weeks post-surgery, mice were given an i.p. injection of 4OHT (20 mg/kg) for three consecutive days directly prior to the onset of the dark cycle to induce recombination of Cre in promoter IV-expressing vHC neurons. 4OHT was prepared by first dissolving 10 mg powder (Sigma, catalog # H6278) in 250 µl of DMSO, and then mixing the 4OHT solution with 400 µl of 25% Tween-80 in 4.35 mL of filtered 1x PBS, for a final 4OHT concentration of 2 mg/ml (Ye et al., 2016). Behavioral testing/sacrifice for immunohistochemistry/*in situ* hybridization was completed 2 weeks post-4OHT injection.

### Immunohistochemistry/Cresyl Violet Staining

For c-Fos immunohistochemistry, mice were trans-cardially perfused with 4% paraformaldehyde 2 h following the termination of tone/context recall. Brains were extracted and stored in 4% paraformaldehyde for 24 h, and then placed into 30% sucrose in 1x PBS/sodium azide (0.05%) for 2-3 d. Coronal sections (50 μm) of the vHC were cut on a sliding microtome (Leica) with attached freezing stage (Physitemp), washed in 5% Tween-80 in 1x PBS, and incubated in blocking solution (0.5% Tween-80, 5% normal goat serum in 1x PBS) with agitation for 6-8 h. The sections were then incubated in 1:1000 anti-Fos antibody (Millipore) in blocking solution overnight at 4°C with agitation. The following day, the sections were washed, incubated in 1:1000 goat anti-rabbit AlexaFluor 647 (Sigma) in blocking solution for 2 h with agitation, washed again, and incubated in 1:5000 DAPI (Sigma) in 1x PBS for 20 min. The sections were then mounted and coverslipped, and co-expression of either tdTomato/c-Fos (for analysis of recruitment of p4-cells during fear behavior) or mCherry/c-Fos (for verification of excitatory DREADD efficacy) was visualized at 20x magnification on a Zeiss 700 LSM confocal microscope. For BDNF immunohistochemistry, mice were trans-cardially perfused with 4% paraformaldehyde/0.15% picric acid/ 0.1% glutaraldehyde in PBS. Brains were cryoprotected in 30% sucrose, then frozen and sectioned (50 μm) and washed in 1x tris-buffered saline (TBS). Slices were incubated in a blocking solution containing 4% normal goat serum/0.1% bovine serum albumin/0.25% Triton-X 100 in TBS for 1 h. Immediately afterward, slices were incubated in mouse on mouse blocking reagent (1:250 in TBS; Vector Labs) for 1 h, washed in TBS, and incubated in 1:200 anti-BDNF antibody (Developmental Studies Hybridoma Bank) at 4°C for 48 h. Following incubation in the primary antibody, slices were again washed in TBS, incubated in 1:1000 goat anti-mouse AlexaFluor 647 (Sigma) in blocking solution for 2 h, and counter-stained with DAPI, mounted, and visualized according to the procedure for c-Fos immunohistochemistry described above (Yang et al., 2014). c-Fos, tdTomato, and BDNF-expressing cells were quantified by eye from maximum intensity projections created from z-stacked, tiled images. For visualization of electrode tracks in the vHC and PrL, we stained microtome-sliced 50 μm coronal sections with 0.1% cresyl violet, and imaged slices with an Olympic BX51 upright light microscope.

### Single-molecule *in situ* hybridization

For verification that exon IV-containing *Bdnf* and *Cre* were co-localized with *tdTomato*, we followed previously published procedures (Colliva et al., 2018). Animals were killed and brains immediately extracted and flash-frozen in 2-methylbutane (ThermoFisher). Coronal sections of vHC (16 µm) were cut, and mounted onto slides (VWR, SuperFrost Plus). The slides were quickly fixed in 10% buffered formalin at 4°C, washed in 1X PBS, and dehydrated in ethanol. Slides were next pre-treated with a protease solution, and subsequently incubated at 40°C for 2 h in a HybEZ oven (ACDBio) with a combination of probes for *Cre, tdTomato,* and exon IV-containing *Bdnf* mRNA. Following incubation with the probes, the slides were incubated at 40°C with a series of fluorescent amplification buffers. DAPI was applied to the slides, and the slides were cover-slipped with Fluoro-Gold (SouthernBiotech). Transcript expression was visualized on a Zeiss LSM 700 confocal microscope with a 40x oil-immersion lens (*Cre*: 555 nm, *tdTomato*: 488 nm, exon IV-containing *Bdnf*: 647 nm, DAPI: 405 nm). For quantification of transcript co-localization, 40x z-stacks from the ventral dentate gyrus were taken, and custom MATLAB functions were used to first isolate cell nuclei from the DAPI channel using the cellsegm toolbox (Hodneland et al., 2013) combined with a watershed algorithm for cell splitting in 3 dimensions. Once centers and boundaries of individual cells were isolated, an intensity threshold was set for transcript detection, and watershed analysis was used to split the remaining pixels in each channel into identified transcripts. Custom MATLAB functions were then used to determine the size of each detected transcript (*regionprops3* function in Image Processing toolbox), and split unusually large areas of fluorescence into multiple transcripts based on known transcript size. Each transcript was then assigned to a cell based on its position in 3 dimensions; transcripts that were detected outside of the boundaries of a cell were excluded from further analysis.

### Behavior

For fear conditioning, animals were briefly handled by the experimenter for 3 days pre-acquisition, and were subsequently conditioned to associate a 30 s tone (4000 Hz) with a 2 s, 0.6 mA foot-shock. 4 tone-shock combinations were given, with the shock co-terminating with the last 2 s of each tone presentation. Tone-shock combinations were separated by 90 s intervals. 48 h later, mice were placed back into the conditioning chamber to measure recall of the conditioning context (in the absence of the tone) for 180 s. Following pre-tone context recall, a total of 23 tones (30 s each) were presented in the absence of the foot-shock for recall of the tone in the fear-associated context (tone/context recall). Tones were separated by 5 s intervals. Freezing (cessation of movement) during acquisition and recall sessions was quantified with automated tracking software (FreezeScan; CleverSys, Inc.). For mice with either hM3Dq or mCherry expression in the vHC, clozapine-N-oxide (CNO; 5 mg/kg, i.p.) dissolved in 1x PBS was administered 45 m. prior to context recall. For mice used to measure c-Fos expression in p4-cells following fear recall, half of the mice were given foot-shocks during acquisition, and half of the mice were presented with the tone in the absence of a foot-shock. In order to assess possible effects of hM3Dq activation on locomotor activity, home-cage activity (movement, rearing) was measured in hM3Dq and mCherry-expressing mice (HomeCageScan software; CleverSys, Inc.) both 1 h prior to and 1 h after CNO injections. We further investigated whether hM3Dq activation caused anxiogenic or anxiolytic effects by administering CNO and measuring time spent in the center and number of center crossings in an open field for 1 h. Position data for open field testing were captured with CaptureStar software (CleverSys, Inc.), and occupancy maps, time spent in the center, and number of center crossings were analyzed with custom MATLAB functions. Fear conditioning, home-cage activity, and open field testing were conducted 7 d apart.

### Electrophysiology and LFP analysis

Local field potentials (LFPs) from the prelimbic (PrL) subregion of the mPFC and the vHC were recorded during context and tone/context recall (Sirenia software; Pinnacle). LFPs were sampled at 2 kHz and bandpass filtered between 0 and 50 Hz. Raw traces were visually inspected for noise artifacts, and de-trended with custom MATLAB functions. Multitaper spectral analysis was used for power density estimation and phase coherence via Chronux toolbox routines in MATLAB (http://chronux.org; Bokil et al., 2010). For phase-amplitude coupling analysis, phase and amplitude values for frequency pairs were extracted via Morlet wavelet convolution. Phase and amplitude values were then binned (num. bins = 18 per cycle), and mean amplitude values were normalized by dividing each bin value by the summed value over all bins. A modulation index value was then derived by calculating the Kullback-Leibler distance between the observed phase-amplitude distribution and a uniform (null) phase-amplitude distribution (Tort et al., 2010):

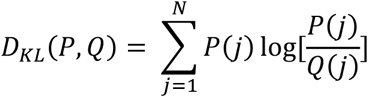

where *P* is equal to the observed distribution, and *Q* is equal to a uniform distribution. For Granger causality, custom MATLAB functions were used to calculate a univariate autoregression of the mPFC LFP at times *t*:

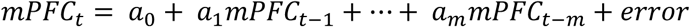

which was augmented with lagged values of the vHC LFP:

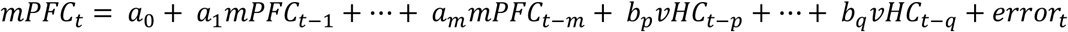

with number of lags being determined by the model order, which was set according to the Bayes’ information criterion (BIC) for each data set. A Granger causal index was then calculated by dividing the index of vHC → mPFC directionality by the total of the indices for both vHC → mPFC and mPFC → vHC directionality. Thus, if the vHC led more strongly, a Granger causal index > 0.5 would be the result, while stronger mPFC directionality would yield a Granger causal index < 0.5.

### Statistical Tests

Individual statistical tests and results of those tests are noted in figure legends throughout the manuscript (statistics performed with GraphPad Prism software). An alpha level of 0.05 was used to determine statistical significance for all tests. For binary classification of *tdTomato* expression based on number of *Bdnf* transcripts, we used logistic regression to predict whether or not *tdTomato* was expressed (cutoff of 5 transcripts) in each cell by training the classifier on transcript data from all cells in each image with the exception of five cells, which were used as a testing set. This procedure was repeated until all cells had been used as test data, and classifier accuracy was calculated by comparing the outcome predicted by the classifier with the training label for each cell. Receiver operating characteristic (ROC) curves were used to validate classifier accuracy, and area under the ROC curve was calculated with custom MATLAB functions. For comparison data, training labels and number of *Bdnf* transcripts were randomly shuffled relative to each other and prediction accuracy and ROC curves were calculated with the shuffled data as described above.

## Acknowledgements

We thank Richard de los Santos Abreu, John Hobbs, Madhavi Tippani, and Danisha Gallop for technical assistance. Funding for these studies was provided by the Lieber Institute for Brain Development and a National Institute of Mental Health R01 to KM (MH105592).

The authors declare no financial, or non-financial, competing interests.

---

**Supplementary Figure 1.**
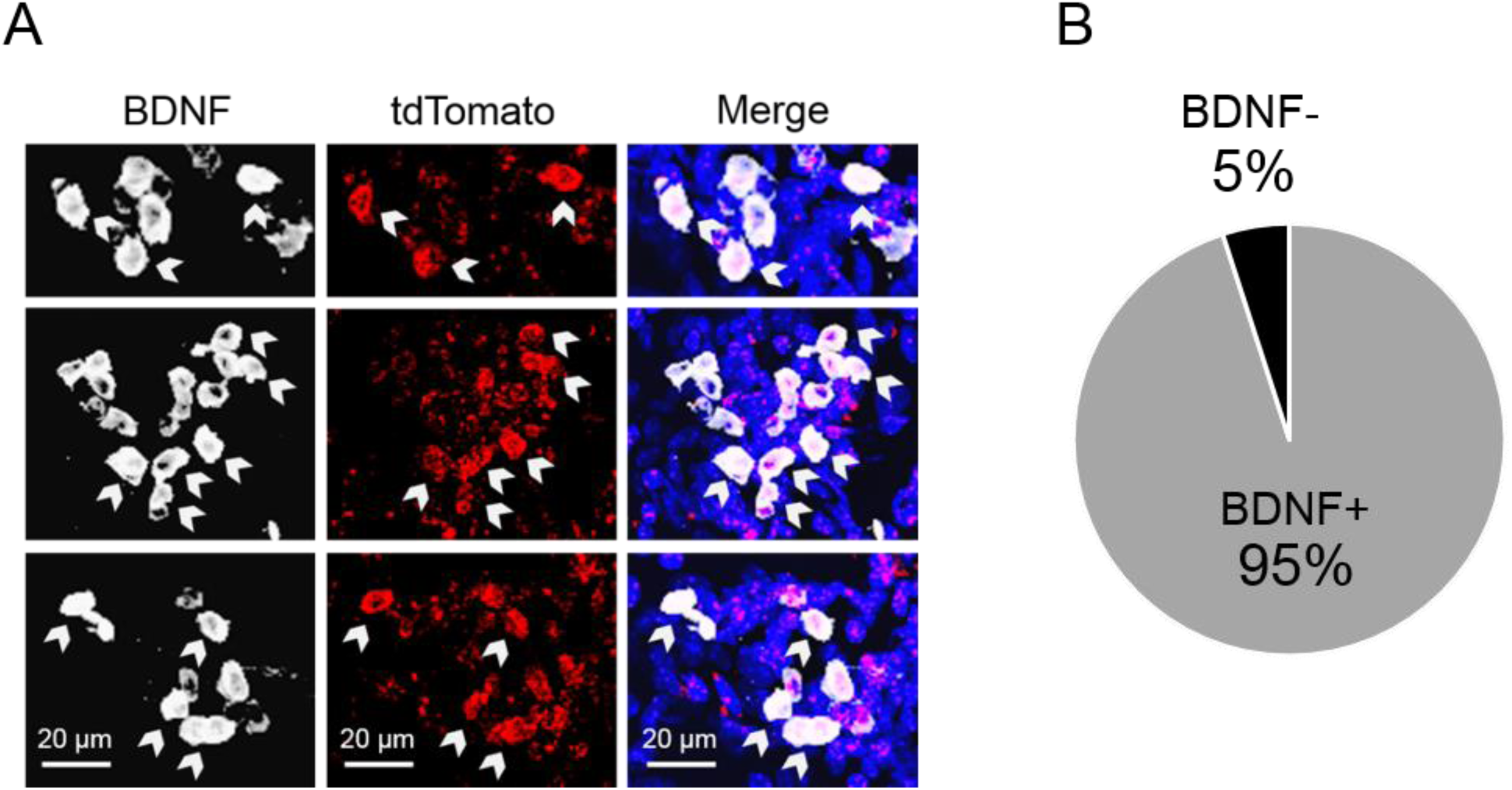
BDNF immunohistochemistry in tdTomato-expressing vHC neurons. *(A)* BDNF immunohistochemistry was used to verify that cells expressing Cre-induced tdTomato (red) also expressed BDNF (white). *(B)* 95% of tdTomato+ cells in the vHC of mice that received injections of both p4Bdnf-ER^T2^-Cre-ER^T2^ and 4OHT co-expressed BDNF protein.

**Supplementary Figure 2.**
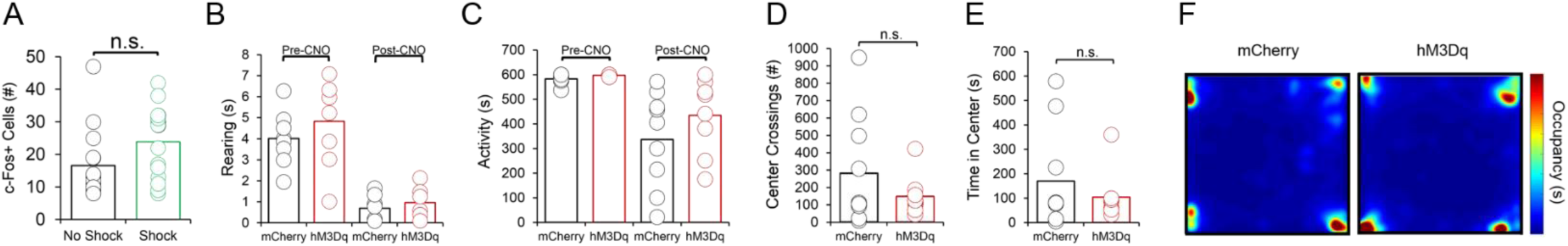
Effects of vHC p4-cell activation on fear recall are fear-specific. *(A)* No significant difference in number of c-Fos+ cells in the vHC following fear recall exists between mice that received a shock during conditioning, and mice that did not receive a shock during conditioning. *(B)* No significant differences in home cage total movement exist between groups either before or after CNO injections. *(C)* No significant differences in rearing in the home cage are observed between groups either before or after CNO injections. *(D)* Activation of vHC p4-cells does not affect number of center crossings, or *(E)* time spent in the center of an open field. *(E)* Open field occupancy does not differ between groups following CNO injections.

**Supplementary Figure 3.**
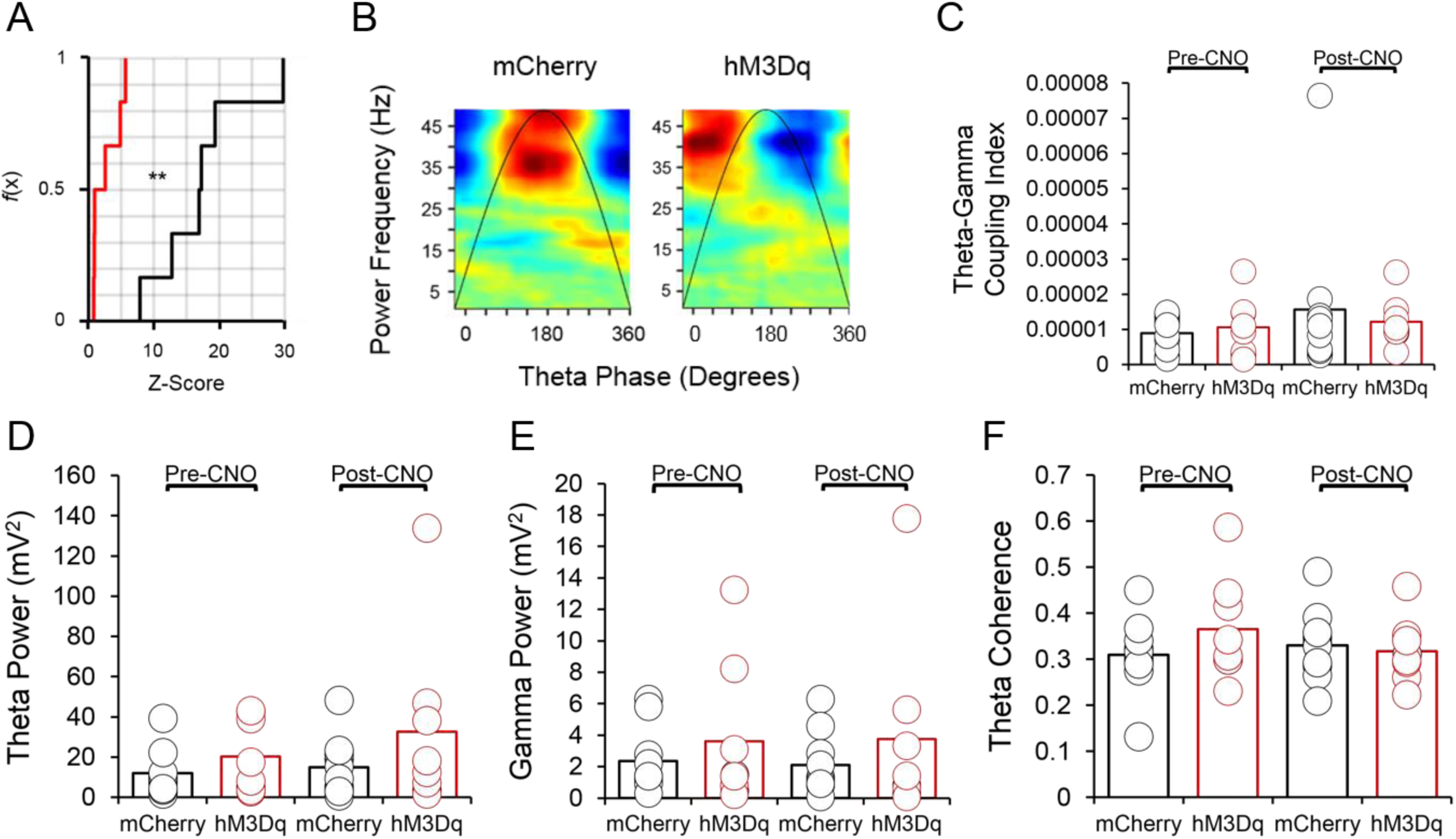
Theta-gamma coupling and LFP in the home cage. *(A)* In order to determine whether observed theta-gamma coupling in the vHC during tone/context recall was greater than what would be expected by chance, phase-amplitude distributions were shuffled 500 times and a theta-gamma coupling index value was obtained for each shuffled distribution. The observed theta-gamma coupling value was compared to the shuffled distribution of coupling values to obtain a z-score. Cumulative frequency distributions of these z-scores were significantly different between hM3Dq (hM3Dq-mCherry in p4-cells) and mCherry (mCherry only in p4-cells) animals (*p* < 0.01, Kolmogorov-Smirnov test), demonstrating that observed theta-gamma coupling following activation of vHC p4-cells was less likely to be higher than chance levels of theta-gamma coupling. *(B)* In addition to a decrease in theta-gamma coupling following CNO injections in hM3Dq mice, slow gamma oscillations shifted their preferred theta coupling phase from the peak to the trough in hM3Dq animals. *(C)* vHC theta-gamma coupling, *(D)* theta power, *(E)* slow gamma power, and *(F)* theta phase coherence between the vHC and PrL did not significantly differ between hM3Dq and mCherry animals in the home cage, either before or after CNO injections (*p* > 0.05 for two-way ANOVAs).

